# Whole genome sequencing identifies high-impact variants in well-known pharmacogenomic genes

**DOI:** 10.1101/368225

**Authors:** Jihoon Choi, Kelan G. Tantisira, Qing Ling Duan

**Author notes:** These authors contributed equally to this work (Kelan G. Tantisira, Qing Ling Duan). Correspondence: QL Duan.

## Abstract

More than 1,100 genetic loci have been correlated with drug response outcomes but disproportionately few have been translated into clinical practice. One explanation for the low rate of clinical implementation is that the majority of associated variants may be in linkage disequilibrium (LD) with the causal variants, which are often elusive. This study aims to identify and characterize likely causal variants within well-established pharmacogenomic genes using next-generation sequencing data from the 1000 Genomes Project. We identified 69,319 genetic variations within 160 pharmacogenomic genes, of which 8,207 variants are in strong LD (r^2^ > 0.8) with known pharmacogenomic variants. Of the latter, 8 are coding or structural variants predicted to have high-impact, with 19 additional missense variants that are predicted to have moderate-impact. In conclusion, we identified putatively functional variants within known pharmacogenomics loci that could account for the association signals and represent the missing causative variants underlying drug response phenotypes.

## Introduction

The current paradigm of drug therapy follows a “trial-and-error” approach where patients are prescribed a drug at a standardized dose with the expectation that alternative therapies or doses will be given during a return clinical visit(s).^1^ Not surprisingly, this is inefficient and potentially hazardous for patients who require urgent care or are susceptible to adverse events, which may result in prolonged suffering and fatalities.^2^ A better understanding of the modulators of drug response will improve and hopefully replace our current trial-and-error approach of drug therapy with more precise methods that are based on scientific knowledge.^3^

To date, more than 1,100 genetic loci have been correlated with drug response phenotypes (The Pharmacogenomics Knowledgebase (PharmGKB): www.pharmgkb.org) but only a small fraction of these genomic findings have been implemented into clinical practice. In 2009, PharmGKB partnered with the Pharmacognomics Research Network (PGRN) to establish the Clinical Pharmacogenetics Implementation Consortium (CPIC)).^4,5,6^ The goal of CPIC is to provide specific guidelines that instruct clinicians on how to use or interpret a patient’s genetic test results to determine the optimal drug and dosage to each patient. As of June 2017, there are 36 drug-gene pairs with CPIC guidelines published, although there are 127 well-established pharmacogenomic genes identified as CPIC genes and 64 additional genes labeled as Very Important Pharmacogenes (VIP) by the PharmGKB curators, which totals to 160 unique genes.

An example of a CPIC guideline is one that instructs physicians on how to interpret genomic information from clinical assays to determine a therapeutic dosage for warfarin, a commonly used drug for the prevention of thrombosis.^7^ Warfarin is known to have a narrow therapeutic index and wide effect variances among patients. For example, a conventional dose of warfarin may not be an effective anticoagulant in some patients or induce adverse events (e.g. excessive bleeding) in others.^8^ Thus, it is often difficult to achieve and maintain a targeted effect by administering conventional doses. Recent advancement in pharmacogenomics helped to facilitate genetic tests of two genes that can be used to predict a patients’ sensitivity to the drug prior to administration. Specifically, the therapeutic dosage of warfarin may be calculated based on one’s genotypes at these loci, which has resulted in a significant improvement in drug safety.^8,9^

Despite the successful translation of a small fraction of pharmacogenomics findings into clinical practice, the rate of clinical implementation has been slow.^6^ One explanation is that the majority of pharmacogenomics loci are correlated with drug response but do not represent the actual, causal variants themselves.^10,11,12^ We hypothesize that the majority of known pharmacogenomics loci are genetic markers that tag causal variants, which have yet to be identified and are likely to be in linkage disequilibrium (LD) with the associated markers. The use of associated variants instead of the causal variants in clinical tests is limiting in that it may not reliably predict drug response.^13^

The primary objective of this study is to identify potentially causal variants in well-established pharmacogenomics-associated genes, which may account for the reported association signals. Specifically, we used whole genome sequencing data from the 1000 Genomes Project^14,15^ to derive all genetic variations identified within the 160 unique CPIC and VIP pharmacogenomics genes. Next, we tested the LD with known pharmacogenomic variants, and determined the predicted function of these LD variants using annotation databases and clinical outcome databases. Our results include a catalog of potentially functional variants that are in LD with well-established pharmacogenomics variants and could represent the causative mutations within these loci.

## Results

### Selection of pharmacogenomics loci and annotation of variants

We selected 127 CPIC genes and 64 VIP genes (total of 160 unique loci) from PharmGKB, which we deemed as “well-established” pharmacogenomics loci **(Supplemental data 1**). Next, we identified 887,980 variants within these loci using next generation sequencing data from the 1000 Genomes Project Phase I, of which 69,319 were variants with minor allele frequencies > 1% **(Supplemental data 2**). Annotation analysis using SnpEff^16^ (genetic variant annotation and effect prediction toolbox) revealed that 65,333 (94%) of these variants were single nucleotide polymorphisms (SNPs), 1,404 (2%) were insertions, and 2,582 (4%) were deletions. As shown in **Figure 1**, the majority of these occur within intronic regions (∼75%), with the remainder located 3’ or downstream (∼11%), 5’ or upstream (∼9%), and exonic (∼2%). Of the coding variants, approximately half of these variants are missense (∼49%), or synonymous mutations (∼50%) with some occurrences of nonsense (∼1%) mutations. We compared our findings with annotation results of whole genome sequencing data of 1000 Genome Project phase I dataset (http://snpeff.sourceforge.net/1kg.html) and confirmed that the results of variant annotation within 160 PGx genes are within an expected range (**Supplemental figure 1**).

**Figure 1.**
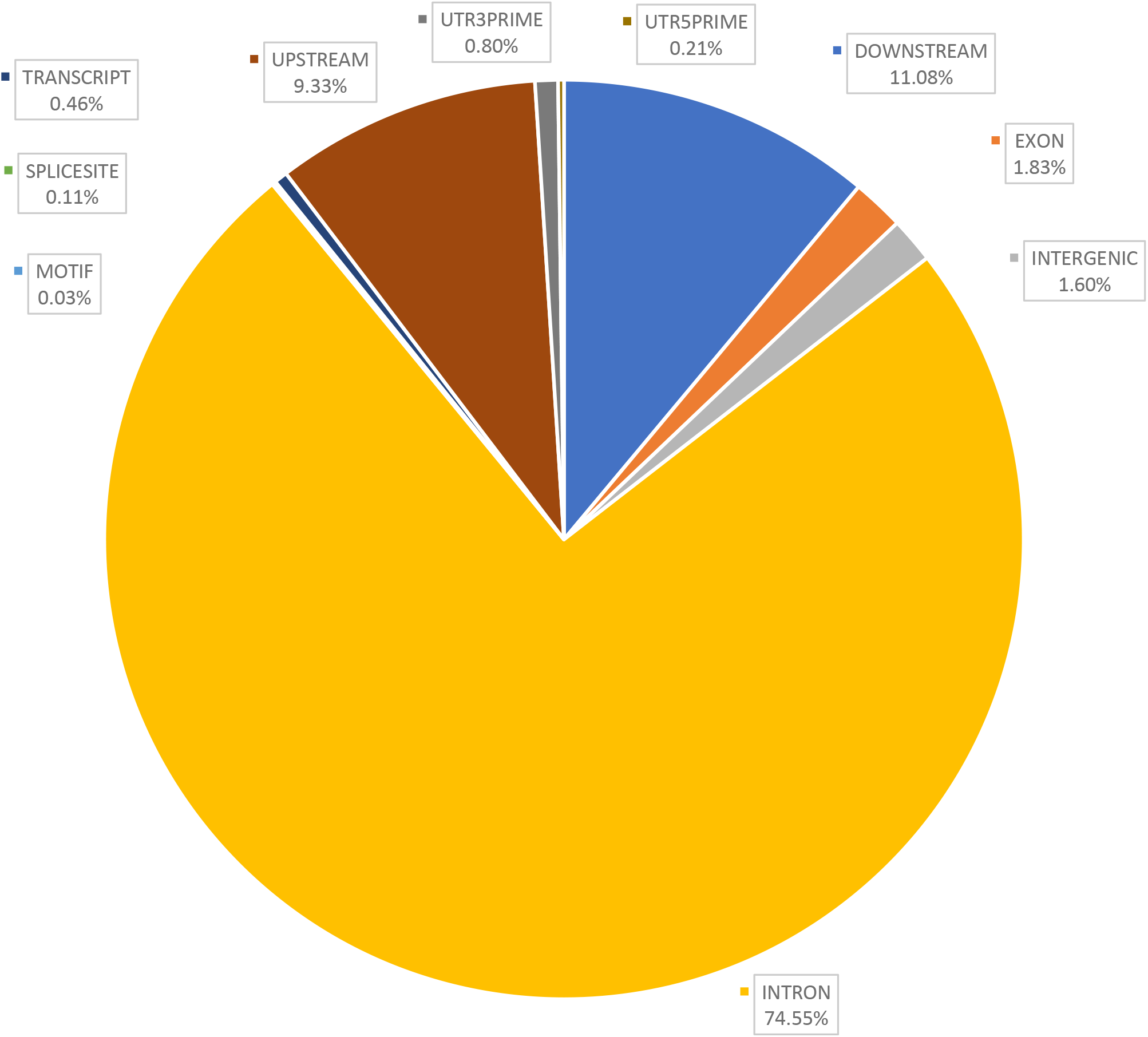
Genomic regions of all variants identified from the 1000 Genomes Project database within 160 known pharmacogenomics genes.

Locations of all the single nucleotide variants identified within the 160 Pharmacogenomics loci using sequence data from the 1000 Genomes Project.

### Linkage disequilibrium analysis

We assessed the LD between associated variants within known pharmacogenomics loci and variants identified in our study. Analysis of LD was done in each of the four populations (American, European, East Asian, African) from Phase I of 1000 Genomes Project. This resulted in 8,207 novel variants forming 21,256 instances of LD (r^2^ > 0.8) with 859 known pharmacogenomics variants (**Supplemental data 3**).

### High-impact variations

We identified 8 variants predicted to have a high-impact using SNPEff from the 1000 GP database that were in LD (r^2^ > 0.8) with 22 known pharmacogenomics variants. These included potentially functional variants that code for an alternative splice donor site, structural interaction, frameshift mutation, stop gain, or stop lost variation. **Table 1** lists these new LD variants along with the corresponding pharmacogenomics variants, the majority of which are predicted to be non-coding located within introns, up/downstream, and synonymous, with only few instances of missense and frameshift variants).

**Table 1.**
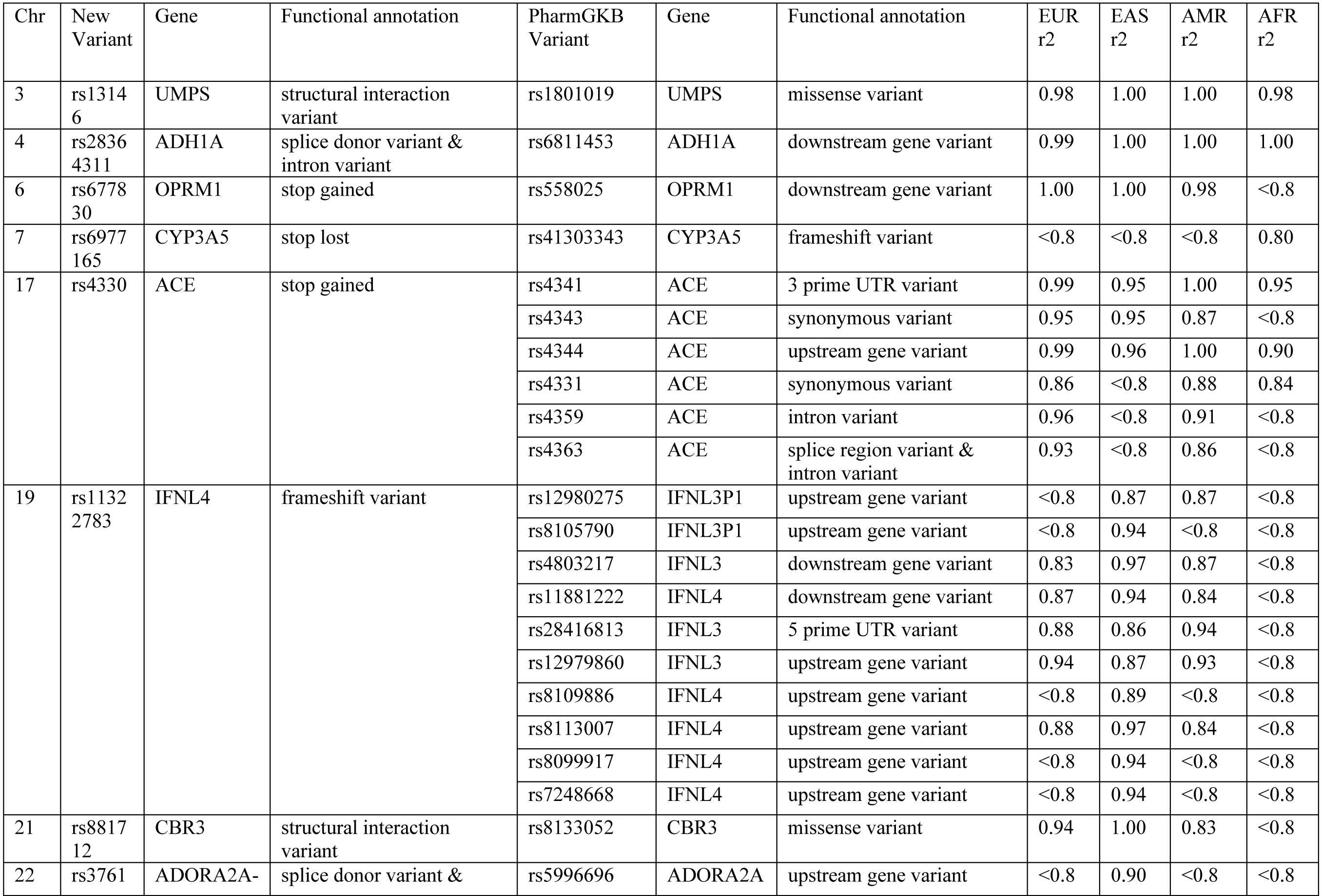
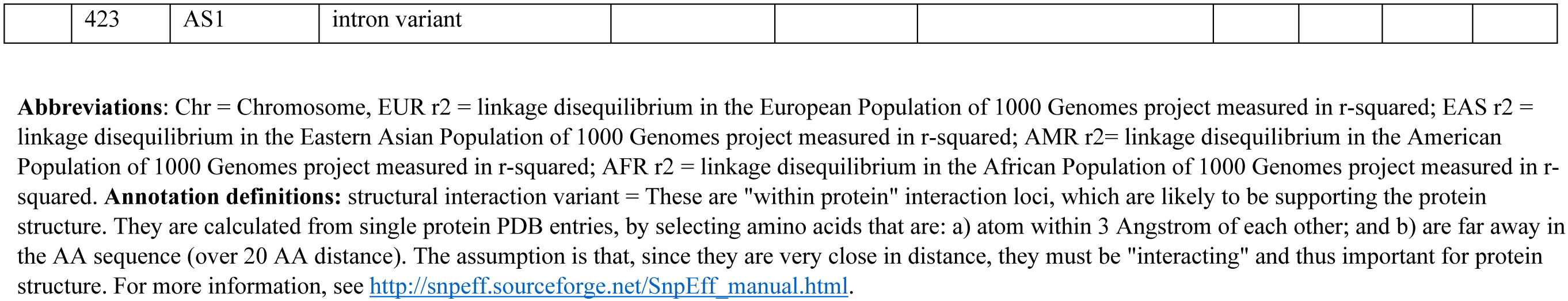
– Variants with high impact predictions, which are in LD with known pharmacogenomics variants.

### Moderate-impact variations

We identified 19 missense variants that are in LD with 32 pharmacogenomics variants, which are predicted to have a moderate, low, or modifying effects by SNPEff (**Table 2)**. Among the newly identified variants, two are regulatory variants that could potentially affect protein binding, and one has been associated with neural tube defects and spina bifida cystica.

**Table 2.** – Variants predicted with moderate impact identified in this study, which are in LD with known pharmacogenomics variants.

| Chr | New Variant | Gene | Functional annotation | PharmGKBVariant | Gene | Annotation | EUR r2 | EAS r2 | AMR r2 | AFR r2 |
| --- | --- | --- | --- | --- | --- | --- | --- | --- | --- | --- |
| 18 | rs2853533<br>★ | C18orf56 | missense variant & TFBS variant | rs2853741 | RP11-806L2.5 | upstream gene variant | <0.8 | 0.85 | <0.8 | <0.8 |
| 1 | rs55867221 | C1orf167 | missense variant & TFBS variant | rs17367504 | CLCN6 | upstream gene variant | <0.8 | 0.9 | <0.8 | <0.8 |
|  |  |  |  | rs3737967 | C1orf167 | missense variant | <0.8 | 0.98 | 0.87 | <0.8 |
|  |  |  |  | rs2274976 | MTHFR | missense variant | <0.8 | 0.96 | 0.87 | <0.8 |
| 1 | rs1537514 | C1orf167 | missense variant | rs3737967 | C1orf167 | missense variant | <0.8 | 0.98 | 0.87 | <0.8 |
|  |  |  |  | rs2274976 | MTHFR | missense variant | <0.8 | 0.96 | 0.87 | <0.8 |
|  |  |  |  | rs17367504 | CLCN6 | upstream gene variant | <0.8 | 0.9 | <0.8 | <0.8 |
| 1 | rs1800595 | F5 | missense variant | rs6018 | F5 | missense variant | 1 | 1 | 1 | 1 |
| 1 | rs6027 | F5 | missense variant | rs6018 | F5 | missense variant | 0.94 | 0.89 | 0.97 | <0.8 |
| 1 | rs6033 | F5 | missense variant | rs6018 | F5 | missense variant | <0.8 | 0.83 | <0.8 | <0.8 |
| 3 | rs3732765 | MED12L | missense variant | rs9859538 | MED12L | intron variant | <0.8 | 0.97 | <0.8 | <0.8 |
|  |  |  |  | rs10935842 | P2RY12 | upstream gene variant | 1 | 0.99 | 0.97 | <0.8 |
|  |  |  |  | rs6798637 | P2RY12 | upstream gene variant | 0.89 | <0.8 | <0.8 | <0.8 |
| 4 | rs1693482 | ADH1C | missense variant | rs1662060 | ADH1C | downstream gene variant | 1 | 1 | 0.96 | 1 |
|  |  |  |  | rs698 | ADH1C | missense variant | 1 | 1 | 0.96 | 1 |
| 4 | rs4963 | ADD1 | missense variant | rs4961 | ADD1 | missense variant | 0.88 | 0.99 | 0.96 | <0.8 |
| 7 | rs2307040 | CALU | missense variant | rs1043550 | CALU | 3 prime UTR variant | 0.82 | <0.8 | 0.96 | 0.89 |
|  |  |  |  | rs11653 | CALU | 3 prime UTR variant | 0.82 | <0.8 | 0.96 | 0.89 |
| 9 | rs56350726 | SLC28A3 | missense variant | rs10868138 | SLC28A3 | missense variant | 0.81 | <0.8 | 0.83 | <0.8 |
| 11 | rs11604671 | ANKK1 | missense variant | rs2734849 | ANKK1 | missense variant | 0.97 | 1 | 0.98 | <0.8 |
|  |  |  |  | rs6277 | DRD2 | synonymous variant | <0.8 | 1 | 0.88 | <0.8 |
|  |  |  |  | rs2587548 | DRD2 | upstream gene variant | <0.8 | 1 | <0.8 | <0.8 |
|  |  |  |  | rs2734833 | DRD2 | upstream gene variant | <0.8 | 1 | <0.8 | <0.8 |
|  |  |  |  | rs1076563 | DRD2 | upstream gene variant | <0.8 | 0.97 | <0.8 | <0.8 |
| 16 | rs115629050 | CES1 | missense variant | rs2307240 | CES1 | missense variant | <0.8 | <0.8 | 0.9 | <0.8 |
| 16 | rs2307227 | CES1 | missense variant | rs2307240 | CES1 | missense variant | <0.8 | <0.8 | 0.9 | <0.8 |
| 16 | rs79711700 | CES1 | missense variant | rs2307240 | CES1 | missense variant | 0.88 | <0.8 | 1 | <0.8 |
| 19 | rs2336219 | CD3EAP | missense variant | rs967591 | CD3EAP | 5 prime UTR variant | 0.83 | 1 | 0.96 | <0.8 |
|  |  |  |  | rs735482 | CD3EAP | missense variant | 1 | 1 | 0.96 | 0.93 |
| 19 | rs12971396 | IFNL4 | missense variant | rs12980275 | IFNL3P1 | upstream gene variant | <0.8 | 0.84 | <0.8 | <0.8 |
|  |  |  |  | rs8105790 | IFNL3P1 | upstream gene variant | 0.92 | 0.97 | 0.97 | <0.8 |
|  |  |  |  | rs4803217 | IFNL3 | downstream gene variant | <0.8 | 0.94 | <0.8 | <0.8 |
|  |  |  |  | rs11881222 | IFNL4 | downstream gene variant | <0.8 | 0.91 | <0.8 | <0.8 |
|  |  |  |  | rs28416813 | IFNL3 | 5 prime UTR variant | <0.8 | 0.83 | <0.8 | <0.8 |
|  |  |  |  | rs12979860 | IFNL3 | upstream gene variant | <0.8 | 0.84 | <0.8 | <0.8 |
|  |  |  |  | rs8109886 | IFNL4 | upstream gene variant | <0.8 | 0.86 | <0.8 | <0.8 |
|  |  |  |  | rs8113007 | IFNL4 | upstream gene variant | <0.8 | 0.94 | <0.8 | <0.8 |
|  |  |  |  | rs8099917 | IFNL4 | upstream gene variant | 0.93 | 0.97 | 0.86 | <0.8 |
|  |  |  |  | rs7248668 | IFNL4 | upstream gene variant | 0.93 | 0.97 | 0.86 | <0.8 |
| 19 | rs4803221 | IFNL4 | missense variant | rs12980275 | IFNL3P1 | upstream gene variant | <0.8 | 0.84 | <0.8 | <0.8 |
|  |  |  |  | rs8105790 | IFNL3P1 | upstream gene variant | 0.93 | 0.97 | 0.95 | 0.81 |
|  |  |  |  | rs4803217 | IFNL3 | downstream gene variant | <0.8 | 0.94 | <0.8 | <0.8 |
|  |  |  |  | rs11881222 | IFNL4 | downstream gene variant | <0.8 | 0.91 | <0.8 | <0.8 |
|  |  |  |  | rs28416813 | IFNL3 | 5 prime UTR variant | <0.8 | 0.83 | <0.8 | <0.8 |
|  |  |  |  | rs12979860 | IFNL3 | upstream gene variant | <0.8 | 0.84 | <0.8 | <0.8 |
|  |  |  |  | rs8109886 | IFNL4 | upstream gene variant | <0.8 | 0.86 | <0.8 | <0.8 |
|  |  |  |  | rs8113007 | IFNL4 | upstream gene variant | <0.8 | 0.94 | <0.8 | <0.8 |
|  |  |  |  | rs8099917 | IFNL4 | upstream gene variant | 0.95 | 0.97 | 0.89 | <0.8 |
|  |  |  |  | rs7248668 | IFNL4 | upstream gene variant | 0.95 | 0.97 | 0.89 | <0.8 |
| 19 | rs762562 | CD3EAP | missense variant | rs967591 | CD3EAP | 5 prime UTR variant | 0.83 | 1 | 0.92 | <0.8 |
|  |  |  |  | rs735482 | CD3EAP | missense variant | 1 | 1 | 1 | 1 |

**rs2853533★** - phenotype association (SNPedia): Neural Tube Defects & Spina Bifida Cystica (The G variant of rs2853533 was associated with Spina Bifida in a transmission disequilibrium test. Study size: 610 families (329 trios, 281 duos) Study population/ethnicity: Patients affected with Spina Bifida and their parents;
Houston, TX; Los Angeles, CA; Toronto, ON, Canada Significance metric(s): $p=0.0213$ ). **Abbreviations:** Chr = Chromosome, EUR $r^2$ = linkage disequilibrium in the European Population of 1000 Genomes project measured in r-squared; EAS $r^2$ = linkage disequilibrium in the Eastern Asian Population of 1000 Genomes project measured in r-squared; AMR $r^2$ = linkage disequilibrium in the American Population of 1000 Genomes project measured in r-squared; AFR $r^2$ = linkage disequilibrium in the African Population of 1000 Genomes project measured in r-squared.

### Low-impact variations

From the total of 8,207 variants in LD, 7,751 variants are classified by SNPEff as variants with unpredictable impact or “modifier” variants. These are in LD with 920 known pharmacogenomics variants with similar impact features. Of these, 324 modifier variants were potential regulatory variants affecting gene expression, protein binding, or transcription factor binding.

In this study, we will focus on modifier variants that are classified under category 1 of RegulomeDB database, which are known eQTLs or variants correlated with variable gene expression. Among 324 modifier variants with RegulomeDB scores, 84 variants were classified as category 1, forming 213 instances of LD with 73 pharmacogenomics variants which are predicted to have low or modifying effects (**Supplemental data 4**).

### Variants associated with clinical outcomes

Using SNPedia database, we discovered 46 variants in LD that are correlated with clinical phenotypes as documented in **Supplemental data 5**.

## Discussion

This manuscript reports the identification of potentially functional genetic variants within genes previously correlated with drug response outcomes. We show that some of the novel variants identified from next-generation sequencing (NGS) of whole genomes (Phase I of the 1000 Genomes Project) are in LD with well-known pharmacogenomics variants and could account for the functional basis underlying the association signals. Many of these LD variants code for non-synonymous amino acid substitutions, frame-shift mutations, introduce a splice variant that results in alternative splicing of the transcript, or located in non-coding regions but are correlated with gene expression levels (expression quantitative trait loci or eQTL) or other clinical phenotypes.

In this study, we used LD analysis to determine the correlation between novel genetic variants identified from the 1000 Genomes Project database and known pharmacogenomics variants. We reasoned that any variant(s) in strong LD (r^2^ > 0.8) with the known pharmacogenomics loci could account for the association signal and have potential to be the actual causal variants at these genomic loci. In order to prioritize the identified variants, we used a popular annotation toolbox (SNPEff) to predict the function of each variant. In addition, we used additional information such as RegulomeDB and SNPedia to prioritize the variant(s) of higher impact from those with low impact.

Many of the variants we identified are “novel” in that these have not been reported in earlier pharmacogenomics studies. For example, we identified a splice donor variant (rs28364311) located on a VIP gene *ADH1A*. This variant is in LD with a pharmacogenomics associated variant, rs6811453, which is associated with increased resistance to cytarabine, fludarabine, gemtuzumab ozogamicin and idarubicin in patients with acute myeloid leukemia.^17^ The associated pharmacogenomics variant is non-coding and have no known biological function as it is located downstream (3’) of the gene. Considering the potential impact of rs28364311 on splicing and its strong LD with the associated pharmacogenomics variant, it is plausible that the splice variant identified is the functional variant that accounts for the original association signals at this locus.

Moreover, we identified that a stop gain variant rs4330 from the VIP gene *ACE*, encoding the angiotensin-converting enzyme, is in LD with 6 known pharmacogenomics variants (rs4341, rs4344, rs4331, rs4359, rs4363, and rs4343). Whereas the latter are intronic or code for synonymous changes, which are less likely to have detrimental effects on the gene product, the identified rs4330 codes for a truncated protein that is likely to have detrimental effects.

Another example is a modifier variant (rs2854509), which we report to be in LD with a pharmacogenomics variant (rs3213239) that is associated with decreased overall survival and progression-free survival when treated with Platinum compounds in patients with non-small-cell lung carcinoma. Our identified variant rs2854509 is located at downstream, whereas pharmacogenomics variant rs3213239 is located upstream of gene encoding X-Ray Repair Cross Complementing 1 protein (*XRCC1*). Our analysis revealed that variant rs2854509 is a cis-eQTL variant acting on CPIC gene *XRCC1*, which is associated with variable efficacy in in platinum-based chemotherapy agents. Additional findings from RegulomeDB showed a direct evidence of binding-site alteration through ChIP-seq and DNase with a matched position weight matrix to the ChIP-seq factor and a DNase footprint. These findings suggest the possibility that rs2854509 has regulatory effects on the gene *XRCC1*, which could modulate response to platinum based chemotherapy treatments.

Our proof of principle study demonstrates that many of the well-known pharmacogenomics loci from PharmGKB are genetic markers that may tag causal variants. Often the latter remain elusive and are likely to be in linkage disequilibrium (LD) with the associated markers. Using NGS data, we identified a number of sequence variants in LD with these pharmacogenomics loci with supporting functional evidence from current annotation softwares. These findings, pending experimental evidence, will ultimately facilitate the translation of improved clinical assays to predict response for a particular drug or dosage prior to administration. The implementation of these clinical tests promises to improve efficacy of drug therapy while reducing the incidence of adverse events.^18^

One limitation of the approach taken is the exclusion of rare variants (minor allele frequency < 0.01). While rare variants are more likely to be functional and clinically relevant, our decision to exclude them from this study was based on the limited sample size (approx. 200-400 in each of the four main populations: American, European, East Asian, African) of 1KGP Phase 1. Specifically, we would not be able to determine LD among rare variants (MAF < 0.01) in such small populations. Another limitation is that this study was based on bioinformatics methods and we did not experimentally validate the potentially functional variants identified, nor confirm their correlation with drug response outcomes. Instead our study was proof of concept that associated variants in well-established pharmacogenomics genes could represent markers of drug response rather than the casual variants. Further studies are needed to identify and ultimately validate the often elusive functional variants in these loci. These additional studies include genotyping of these potentially functional variants (identified in LD with the associated variants) and testing them directly for correlation with drug response outcomes in clinical trials. Other experiments are needed to confirm the biological impact of these variants on the resultant RNA transcripts or proteins, which depends on the predicted impact of the variants identified. For example, variants of high impact (Table 1) include splicing effects, premature stop codons, and structural interactions, which could be validated through direct sequencing of transcripts and mass spectrometry to detect truncated and mis-folded proteins.

Our study identified novel genetic variations located in well-established pharmacogenomics genes, which could account for the association signals at these loci and have strong impact on the resulting gene products. We applied an innovative approach that combined bioinformatics resources such as PharmGKB, sequencing data from the 1000 GP, population annotation software such as SNPEff as well as databases such as RegulomeDB to identify novel variants and predict their functional effects within pharmacogenomics loci. Moreover, we determined that a number of these potentially functional variants are in LD with known pharmacogenomics variants and could account at least in part for the original association signals. Identification of these elusive causal variants could facilitate more accurate genetic tests to predict treatment response prior to drug administration. The improved accuracy results from direct testing instead of relying on LD, which varies among populations (as noted by our study of LD across 4 populations in the 1000 GP). Thus, identification of causal variants will improve the translation of pharmacogenomics findings into clinical practice and ultimately replace the current trial and error approach for drug therapy, moving us closer towards precision medicine.

## Methods

### Pharmacogenomic genes

We selected 160 unique pharmacogenomics associated loci, containing 127 CPIC genes (June 5^th^, 2017 release) and 64 VIP genes (May 1st, 2017 release) from the PharmGKB database. Then, we identified the genomic coordinates of each gene from the GRCh37/hg19 assembly of the human reference genome using the University of Santa Cruz (UCSC) Genome Browser.^19^ Next, genomic coordinates were padded with 5000 bp both 5’ and 3’ of each gene to include potential regulatory regions. All variants that appear in at least 1% of the 1000 Genomes Project Phase I population (Feb. 2009 release) were extracted.

### Functional annotations

After reviewing many annotation tools (including annoVar, VEP, Polyphen/SIFT, CADD), we decided that SnpEff best meets our needs as it allows a great degree of compatibility with various input formats, offers high flexibility in search settings, can annotate a full exome set in seconds, based on up-to-date transcript and protein databases, and has the ability to be integrated with other tools. SnPEff (version 4.2, build 2015-12-05) was used with the GRCh37.75 assembly to predict the effects of identified variants. For variants with multiple annotations (e.g. variant affects multiple genes or have varying effects depending on the transcript), only the most severe consequence was selected and used to represent each variant in tables to ease the comparison of impacts among variants. To standardize terminology used for assessing sequence changes, SNPEff uses sequence ontology (http://www.sequenceontology.org/) definitions to describe functional annotations.

### Linkage disequilibrium analysis

Linkage disequilibrium (LD) between the well-established pharmacogenomics variants (1,151 variants annotated by PharmGKB retrieved on June 16^th^, 2017, that are found within 160 PGx loci and 1000 Genomes project phase 1 dataset) and identified variants from the 1000 Genomes Project phase 1 dataset using Plink (version 1.09).^20^ Distance window for the LD analysis were set to 1Mb and an r^2^ threshold of > 0.8.

### SNPs associated with regulation and phenotypes

For each variant identified to be in LD with an established pharmacogenomic variant, we used RegulomeDB^21^ to evaluate and score those that have the potential to cause regulatory changes, such as eQTL, regions of DNAase hypersensitivity, binding sites of transcription factors and proteins. RegulomeDB uses GEO^22^, the ENCODE^23^ project, and various published literatures to assess these information. In addition to that, we used SNPedia^24^, a database of over 90,000 SNPs and associated peer-reviewed scientific publications, to identify variants that are previously associated with phenotypes. (**Figure 2**)

**Figure 2.**
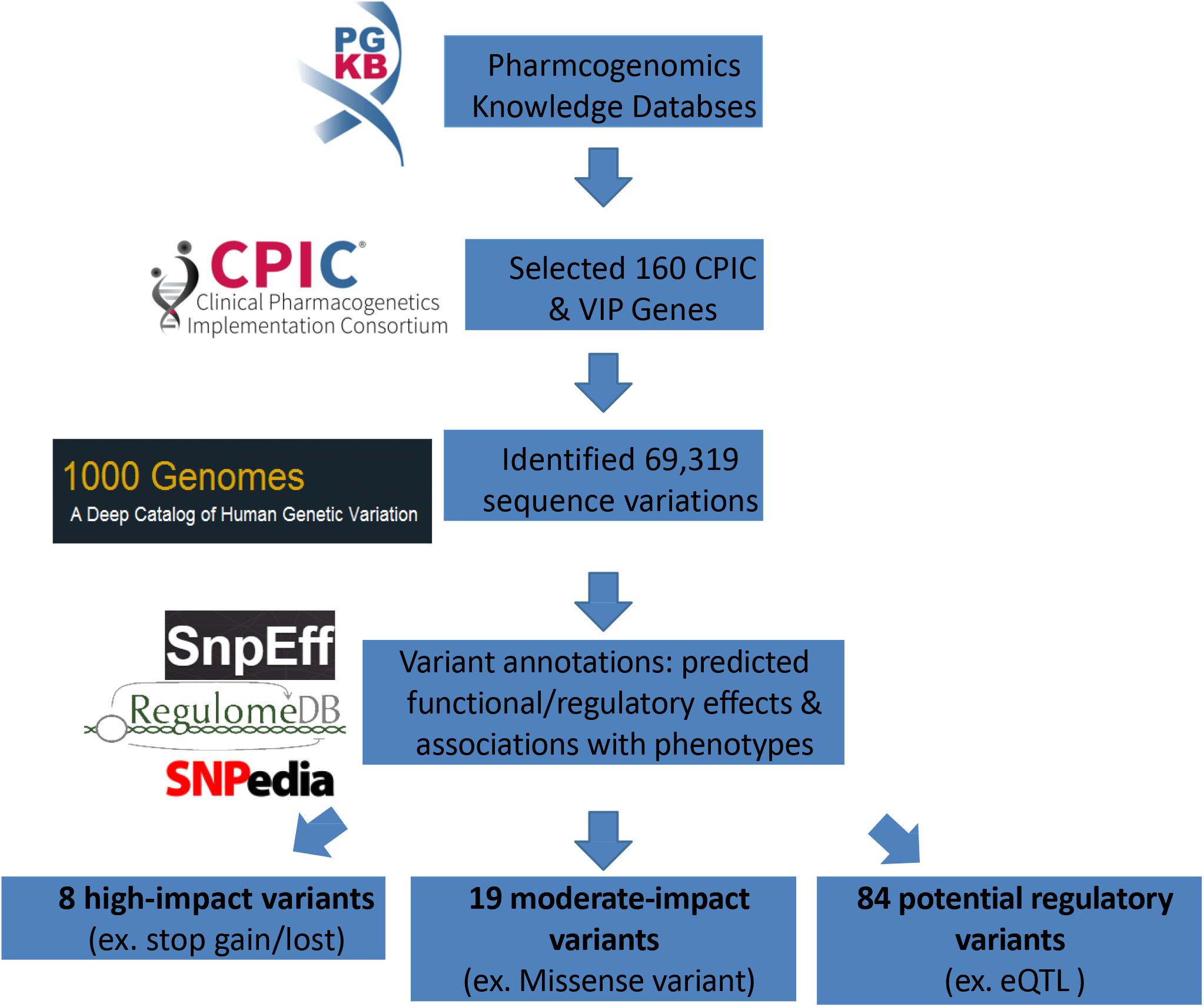
Overview of the experimental design. Flow of work outlined in methods section of the manuscript, which highlights the selection of 160 genes from the Pharmacogenomics Knowledge Database (PharmGKB), identification of variants from the 1000 Genome Project Data, and subsequent steps for annotation and test LD among variants.

## Acknowledgements

We would like to thank Dr. Alan R. Shuldiner and support from the Translational Pharmacogenomics Project. This manuscript was supported by NIH grants U01 HL65899, U01 HL105198 and K99 HL116651. Computations were performed on resources and with support provided by the Centre for Advanced Computing (CAC) at Queen’s University in Kingston, Ontario. The CAC is funded by: the Canada Foundation for Innovation, the Government of Ontario, and Queen’s University. QLD receives funding from the Canadian Institutes of Health Research and Queen’s University.

## Conflict of interest

The authors declare no conflicts of interest.

## Author contribution

All authors contributed to the writing of the manuscript. J.C. performed the data analyses and drafted the manuscript. Q.L.D. supervised data analyses and assisted in the writing of the manuscript. Q.L.D. and K.G.T. designed the research project.

## Code Availability

Code and data used in this manuscript can be accessed from a public repository https://github.com/12jc59/DuanlabPharmacogenomicsProject.

---

**Supplemental Figure 1.**
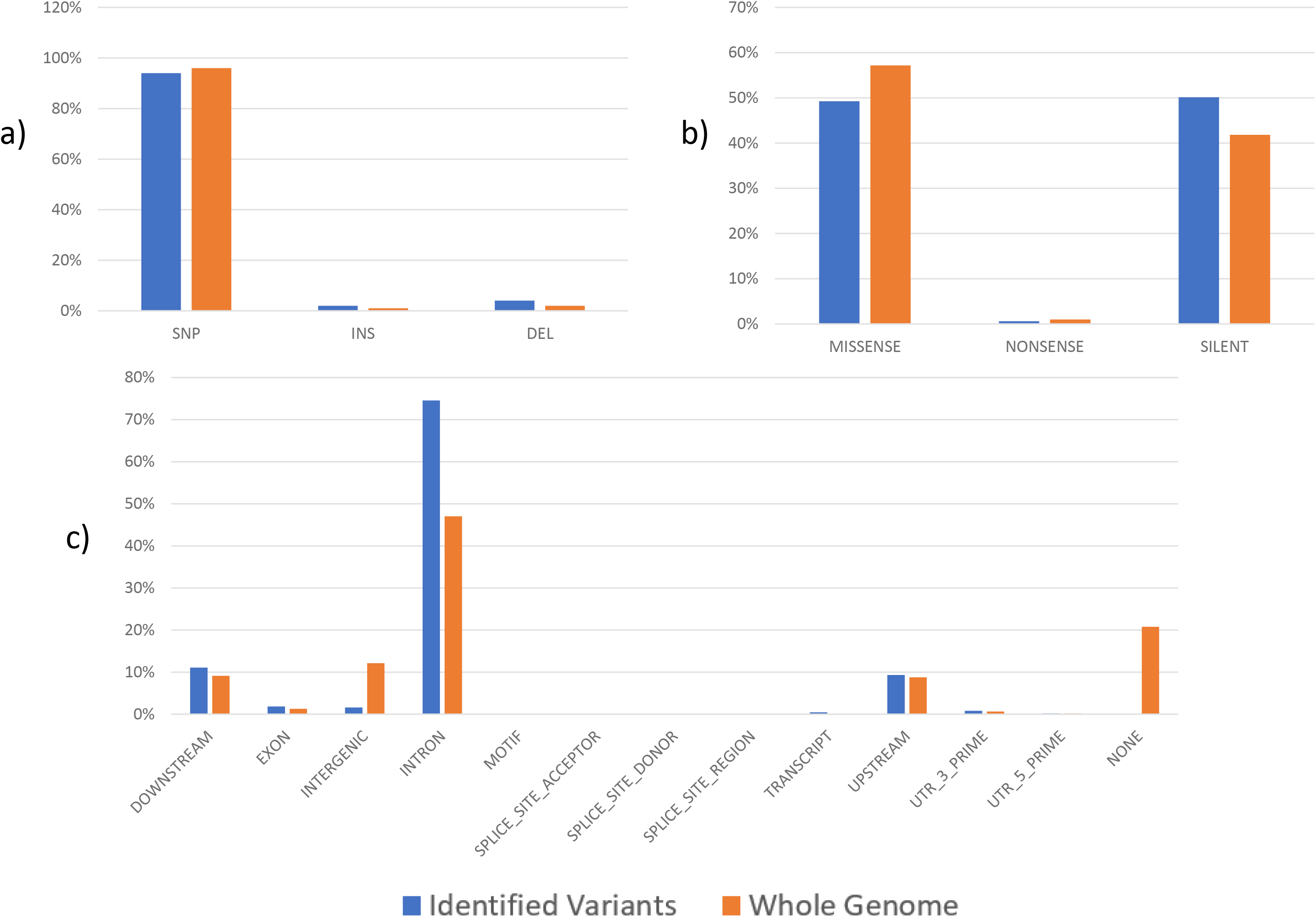
Comparison of annotation findings between variants from 160 PGx genes and whole genome. a) 94% of variants identified in 160 PGx genes were SNPs, 4% deletions, 2% insertions. These numbers are similar to the whole exome data from the 1000 GP. b) Annotation of coding regions within 160 PGx genes identified 49% missense, 50% silent, and 1% nonsense variations. Annotation results from the entire exome had a slightly higher rate of missense mutations and lower rate of silent mutations. However, the ratio of missense to silent mutations in the human exome is expected to be approx. 1.0. Thus, we concluded that our findings fall within the expected range. c) In both whole genome and the 160 PGx genes, the majority of variants fall within intronic regions. Whole genome annotations resulted in higher number of intergenic variants (∼12%) compared 160 PGx genes (∼1.5%). This is due to the fact that we had included limited (5000 bp) flanking regions in our targeted PGx genes in contrast to whole genome sequences. Other than intergenic regions, variants located 5’, 3’, exons, and splice sites occurred with similar frequencies in our candidate genes compared to the whole genome.

## References

1 Evans WE, Relling M V. Moving towards individualized medicine with pharmacogenomics. Nature 2004; 429: 464–468.

2 Giacomini KM, Yee SW, Ratain MJ, Weinshilboum RM, Kamatani N, Nakamura Y. Pharmacogenomics and patient care: one size does not fit all. Sci Transl Med 2012; 4: 153ps18–153ps18.

3 Evans WE, Relling M V. Pharmacogenomics: translating functional genomics into rational therapeutics. Science (80-) 1999; 286: 487–491.

4 Hewett M, Oliver DE, Rubin DL, Easton KL, Stuart JM, Altman RB et al. PharmGKB: the pharmacogenetics knowledge base. Nucleic Acids Res 2002; 30: 163–165.

5 Shuldiner AR, Relling M V, Peterson JF, Hicks K, Freimuth RR, Sadee W et al. The Pharmacogenomics Research Network Translational Pharmacogenetics Program: Overcoming Challenges of Real-World Implementation. Clin Pharmacol Ther 2013; 94: 207–210.

6 Relling M V, Klein TE. CPIC: clinical pharmacogenetics implementation consortium of the pharmacogenomics research network. Clin Pharmacol Ther 2011; 89: 464–467.

7 Johnson JA, Gong L, Whirl-Carrillo M, Gage BF, Scott SA, Stein CM et al. Clinical Pharmacogenetics Implementation Consortium Guidelines for CYP2C9 and VKORC1 genotypes and warfarin dosing. Clin Pharmacol Ther 2011; 90: 625–629.

8 Jaffer A, Bragg L. Practical tips for warfarin dosing and monitoring. Cleve Clin J Med 2003; 70: 361–371.

9 Takeuchi F, McGinnis R, Bourgeois S, Barnes C, Eriksson N, Soranzo N et al. A genome-wide association study confirms VKORC1, CYP2C9, and CYP4F2 as principal genetic determinants of warfarin dose. PLoS Genet 2009; 5: e1000433.

10 Soranzo N, Cavalleri GL, Weale ME, Wood NW, Depondt C, Marguerie R et al. Identifying candidate causal variants responsible for altered activity of the ABCB1 multidrug resistance gene. Genome Res 2004; 14: 1333–1344.

11 Wechsler ME, Israel E. How pharmacogenomics will play a role in the management of asthma. Am J Respir Crit Care Med 2005; 172: 12–18.

12 Zhang W, Dolan ME. Impact of the 1000 genomes project on the next wave of pharmacogenomic discovery. Pharmacogenomics 2010; 11: 249–256.

13 Van den Broeck T, Joniau S, Clinckemalie L, Helsen C, Prekovic S, Spans L et al. The role of single nucleotide polymorphisms in predicting prostate cancer risk and therapeutic decision making. Biomed Res Int 2014; 2014.

14 Whirl-Carrillo M, McDonagh EM, Hebert JM, Gong L, Sangkuhl K, Thorn CF et al. Pharmacogenomics knowledge for personalized medicine. Clin Pharmacol Ther 2012; 92: 414–417.

15 The 1000 Genomes Project Consortium. A global reference for human genetic variation. Nature. 2015; 526: 68–74.

16 Cingolani P, Platts A, Wang le L, Coon M, Nguyen T, Wang L et al. A program for annotating and predicting the effects of single nucleotide polymorphisms, SnpEff: SNPs in the genome of Drosophila melanogaster strain w1118; iso-2; iso-3. Fly 2012; 6: 80–92.

17 Iacobucci I, Lonetti A, Candoni A, Sazzini M, Papayannidis C, Formica S et al. Profiling of drug-metabolizing enzymes/transporters in CD33+ acute myeloid leukemia patients treated with Gemtuzumab-Ozogamicin and Fludarabine, Cytarabine and Idarubicin. Pharmacogenomics J 2013; 13: 335–341.

18 Mancinelli L, Cronin M, Sadée W. Pharmacogenomics: the promise of personalized medicine. AAPS J 2000; 2: 29–41.

19 Karolchik D, Hinrichs AS, Furey TS, Roskin KM, Sugnet CW, Haussler D et al. The UCSC Table Browser data retrieval tool. Nucleic Acids Res 2004; 32: D493–6.

20 Purcell S, Neale B, Todd-Brown K, Thomas L, Ferreira MAR, Bender D et al. PLINK: A tool set for whole-genome association and population-based linkage analyses. Am J Hum Genet 2007; 81: 559–575.

21 Boyle AP, Hong EL, Hariharan M, Cheng Y, Schaub MA, Kasowski M et al. Annotation of functional variation in personal genomes using RegulomeDB. Genome Res 2012; 22: 1790–1797.

22 Edgar R. Gene Expression Omnibus: NCBI gene expression and hybridization array data repository. Nucleic Acids Res 2002; 30: 207–210.

23 Consortium EP, Dunham I, Kundaje A, Aldred SF, Collins PJ, Davis C a et al. An integrated encyclopedia of DNA elements in the human genome. Nature 2012; 489: 57–74.

24 Cariaso M, Lennon G. SNPedia: a wiki supporting personal genome annotation, interpretation and analysis. Nucleic Acids Res 2012; 40: D1308–12.

